# *DFNA5*-mediated pyroptosis is a driver for venetoclax and azacytidine synergy in myeloid leukemia

**DOI:** 10.1101/2023.11.17.567421

**Authors:** Aarthi Nivasini Mahesh, Joanne Lai Xin-Yi, Mandy Tng Jia Lin, Weilin Lin, Jia Cheng Wan, Junho Yoon, Kaiwen Chen, Shruti Bhatt

## Abstract

Acute myeloid leukemia (AML) remains the deadliest adult leukemia with dismal clinical outcomes. Since 2020, a combination of BCL-2 inhibitor (venetoclax, VEN) with hypomethylating agent (azacytidine/decitabine, AZA/DAC) has become a new standard of care in elderly or unfit AML patients. However, the underlying mechanism of synergy between venetoclax and azacytidine combination is not well understood. While apoptosis is regarded as the primary mode of cell death mechanism caused by the venetoclax and azacytidine combination, we provide novel evidence for pyroptosis as additional cell death mechanisms in response to venetoclax and azacytidine combination therapy. We found that long-term treatment with azacytidine caused hypomethylation and significant upregulation in DFNA5/GSDME, pore forming Gasdermin family gene that is otherwise silent in myeloid leukemia. We found that azacytidine mediates N-terminal pore-forming DFNA5 cleavage, membrane rupture, and subsequent pyroptosis of *DFNA5* overexpressing cells in response to venetoclax and azacytidine. Deletion of *DFNA5* reduced total cell viability, where *DFNA5* KO cells exclusively underwent apoptosis while DFNA5 OE cells showed increased propidium iodide uptake, a marker for membrane rupture. Overall, our study establishes *DFNA5* as an important mediator of venetoclax and azacytidine-induced cell death via non-apoptotic mechanisms.

## Introduction

Acute Myeloid Leukemia (AML) is the most common form of leukemia in adults with dismal outcomes. AML originates from the uncontrolled proliferation and accumulation of immature myeloblast or myeloid progenitor cells in the bone marrow. The rapid proliferation and accumulation of immature myeloblasts impairs normal production of monocytes, platelets, neutrophils, white blood cells (WBC) and red blood cells (RBC) leading to anaemia, thrombocytopenia, and several infections [1]. The standard induction regimen (7+3 induction) for AML patients is a combination of cytarabine administered for 7 days followed by daunorubicin for 3 days (7+3 regimen) [2]. After one or two rounds of induction therapy, 75% of younger adults <50 years and 50% of the adults >60 years achieve complete remission (CR) [3]. The 5-year survival rate is 57.1% for the patients with age <50 years and 7.5% for > 65 years old [4].

Almost half of the patients, diagnosed with AML, are >75y/o-attain poor tolerance and response rate to intensive 7+3 chemotherapy regimen [5]. Additionally, daunorubicin used in the combination therapy has high cardiotoxicity in elderly patients [6]. In such cases, less intensive therapies, or monotherapies such as hypomethylating agents (HMAs) or low dose cytarabine (LDAC) were tested. However, this method did not improve the response rate (RR), nor did it prolong the survival in elderly patients [7]. In April 2016, BCL-2 inhibitor venetoclax (VEN) was approved for CLL patients with 17p chromosomal deletion. venetoclax as a monotherapy in AML showed limited clinical response of 19% in relapse or refractory (R/R) AML [8]. However, in combination with HMA, in treatment naïve elderly AML, venetoclax showed impressive clinical response of 73% (CR+CRi) resulting in clinical approval of venetoclax in combination with Azacytidine (AZA) or decitabine (DAC) and Low dose cytarabine (LDAC) for >75y/o AML patients that are unable to undergo intensive chemotherapy [9,10] and younger patients with two or more co-morbidities [11,12].

venetoclax selectively kills BCL-2 dependent tumor cells via inducing mitochondrial pathway of apoptosis [13]. venetoclax binds to BCL-2 by displacing the BCL-2 bound pro-apoptotic protein BIM. BIM then activates pro-apoptotic BAX and BAK to restore the apoptosis in cancer cell [14,15]. Azacytidine and decitabine are hypomethylating agents that inhibits DNA methyl transferase (DNMT). Although the precise identification of target genes remains unclear, downregulation in the expression of several tumour suppressor genes is often reported as primary mode of tumor growth across many cancer types [16]. Additionally, azacytidine upregulates expression of genes involved in tumor suppression, apoptosis, cell cycle, and DNA repair mechanisms, including cyclin-dependent kinase inhibitor 2B (CDKN2B), tissue inhibitor of metalloproteinase 3 (TIMP3), p16, cyclin-dependent kinase inhibitor 1C (CDKN1C), and RAS association domain family 1 (RASSF1)[17]. However, the precise mechanism by which azacytidine induces anti-leukemic effects in myeloid malignancies such as AML and myelodysplastic syndrome (MDS) remain unclear [18].

In 2018, two studies proposed that independent of its demethylation property, azacytidine increases the expression of PUMA and NOXA (BH3-only protein) in AML cells by inducing integrated stress response pathway (ISR) [19,20]. Thus, AML cells are sensitized or primed by azacytidine for venetoclax mediated apoptosis explaining the synergistic killing of the combination therapy [21]. JB Lee et. al. suggested potential role of immune mediated killing by venetoclax and azacytidine combination therapy. This study showed venetoclax activates T cell by increasing the ROS production via inhibition of respiratory chain super complex while azacytidine triggers the cGAS/STING pathway, resulting in viral mimicry. Thus, azacytidine makes the cells more vulnerable to T-cell mediated toxicity activated by venetoclax [22]. However clinical evidence of proposed mechanistic studies is still limited. Collectively, synergistic cytotoxicity and mechanism of action for combination of Azacytidine+venetoclax mediated clinical benefit is still not completely understood.

Besides apoptosis, other forms of programmed cell death such as pyroptosis could also induce cytotoxic response after chemotherapy treatment[24]. Pyroptosis is an inflammatory form of programmed cell death mediated by a group of gasdermin protein superfamily, including *GSDMA, GSDMB, GSDMC, GSDMD*, *GSDME* (or *DFNA5*) and Pejvakin (*PJVK*). The necrotic property of the gasdermins is activated when inflammatory caspases cleave the Gasdermin N-terminal (*GSDM*-NT) from the C-Terminal (*GSDM*-CT) [25]. Cleaved gasdermins undergo oligomerization to form pores in the plasma membrane resulting in membrane rupture [26]. Subsequently, the inflammatory molecules such as IL-18, IL-1β, IL-1α and HMGB1 are released into the extracellular space [27–29]. *GSDME* N-Terminal (*GSDME*-NT/*DFNA5*-NT) and *GSDMD* N-Terminal (*GSDMD*-NT) proteins carry pore forming ability to induce pyroptosis. Other Gasdermins also show pore forming property, but they do not get activated in response to pathological or physiological stimuli. A recent study reported that *DFNA5* has tumour suppressive role in melanoma, breast, colon and cervical carcinoma models [23]. While *DFNA5* is cleaved by caspase 3 activated in response to chemotherapeutic agents like cisplatin, paclitaxel and fluorouracil [30], a number of solid cancers show epigenetic silencing including breast, colorectal and gastric cancer [31–33]. There are also reports showing decitabine increasing the expression of *DFNA5* [31].

In the current study, we aimed to investigate contribution of non-apoptotic forms of cell death in azacytidine+venetoclax combination mediated cytotoxicity in AML cells. We discovered that azacytidine upregulates *DFNA5* by hypomethylation. We found that *DFNA5* is cleaved by azacytidine to form *DFNA5*-NT and subsequently oligomerizes to cause pyroptosis of AML cells. Further, knockout of *DFNA5* prevents pyroptosis and instead primes cells for apoptosis.

## Materials and Methods

### Cell lines and reagents

The AML cell lines MOLM-13, MOLM-14, THP-1, NB4, HL-60, Kasumi-1, MV411, AML-2, AML-3 were cultured in RPMI 1640 medium (Gibco) supplemented with 10% fetal bovine serum (FBS) (Gibco) and 1% penicillin/streptomycin (P/S) (Gibco). All cells were grown at 37 °C in a 5% CO_2_ incubator. Venetoclax (VEN), Z-DEVD-FMK (caspase-3 inhibitor), Q-VD-Oph (pan caspase inhibitor), Azacytidine (AZA), puromycin was purchased from MedChemExpress.

### Plasmids and cloning

Full-length *DFNA5* was expressed in AML cell lines with the help of lentiviral plasmid obtained from addgene #154876. To knock out *DFNA5* we cloned sgRNA sequence obtained from Dr Fang shao lab [24] into the lenti-CRISPR-V2 mCherry plasmid addgene plasmid # 99154. We obtained SgRNA as reverse and forward primer from Integrated DNA technologies and annealed it at 37^0^C for 30min. We next cleaved the Lenti-CRISPR-V2 mCherry with BsmB1-V2 for 1.5hrs at 55^0^C and eluted the fragment of interest from the agarose gel. We then ligated the annealed SgRNA with BsmB1 cleaved Lenti-CRISPR-V2 with T4 DNA ligase at 16^0^C overnight. After transformation we selected 10 colonies and expanded to isolate plasmid with ampicillin selection. Finally, the cloned plasmid was verified for cloning by sequencing.

### Transfection and transduction

HEK293T cells were grown in DMEM with no P/S (Gibco) for 24 hours. *hDFNA5* Lenti-viral transfection was carried out using Polyfect™ (QIAGEN) along with psPAX2 and pMD2G with the ration of 1:1:2. The viral supernatant was collected for 2 days and used for spinfection in AML cell lines (MOLM-14, MOLM-13, Jurkat, AML-2, and THP-1) using 10µg/ml polybrene. The transfection efficiency was checked after (i) 24 hours using FACS for BFP and mCherry positive cells or (ii) puromycin selection for one week and performing western blot analysis.

### Methylation Specific PCR (MSP)

Genomic DNA was isolated from MOLM-13 cell lines after treatment with azacytidine (0.5µM). The genomic DNA was bisulfite converted over night with EZ DNA methylation kit (D5001). Then the bisulphite converted DNA was used for performing Methylation-Specific PCR with methylated sets of primers (F-5’AGGAGGCGTCGTTTTTAAATTTTA 3’, R-5’ ATTACTAACGACCCAACGCTCAA 3’) obtained from [34]. The PCR conditions include 1. Denaturation-95^0^C (2min) 2. Annealing-60^0^C (20sec) 3. Extension-72^0^C (20sec) 4. Final extension-72^0^C (5min) for 40 cycles. Finally, the results were analysed by 2% agarose gel electrophoresis.

### RNA isolation, Reverse transcription, and qPCR

RNA was isolated using the RNeasy^®^ Plus Micro Kit (QIAGEN) and cDNA was prepared as per the GoScript™ Reverse Transcription System (Promega) protocol. qPCR was conducted using AceQ® SYBR green kit (Biobasic). *DFNA5* primers (F-5’ CCAGTTTTTATCCCTCACCCTTG 3’) and R-5’ CAAACTTGCCCTCGTATTTCACA3’) GAPDH primers (F-5’ CATCACTGCCACCCAGAAGACTG and R-5’ ATGCCAGTGAGCTTCCCGTTCAG 3’) were purchased from Integrated DNA technologies and used for RT-qPCR runs, conducted using 96-well PCR plates using Bio-Rad CFX-96 and the gene expression (ΔΔCq) was quantified using Bio-Rad CFX Manager 3.1.

### Immunoblotting

Cell proteins were extracted using RIPA Lysis Buffer (Thermo Fisher) with freshly added protease/phosphatase cocktail inhibitor 100x (Cell Signalling Technology). Protein concentrations were determined using BCA protein assay kit (Thermo Fisher). 30-40 µg of protein with SDS loading dye (4x) was loaded onto 12% polyacrylamide gels. Proteins were transferred onto nitrocellulose membranes using semi dry turbo transfer system. Then the membrane was incubated for overnight with primary antibodies diluted 1:1000 in 5 % BSA in TBST. Blots were washed with PBST (3X) before and after the secondary antibodies prior to imaging. Chemiluminescence images were obtained using ChemiDoc Imaging Systems (Bio-Rad) using supersignal^TM^ west femto maximum sensitivity substrate reagents (#34095). Recombinant Anti-*DFNA5*/*GSDME* antibody – N Terminal (Abcam), Anti-*DFNA5*/*GSDME* antibody – Full length (Abcam), Anti-GAPDH antibody (Abcam).

### Non-reducing PAGE

Cell proteins were extracted using RIPA Lysis Buffer (Thermo Fisher) with freshly added protease/phosphatase cocktail inhibitor 100x (Cell Signalling Technology). Protein concentrations were determined using BCA protein assay kit (Thermo Fisher). 10 µg of protein with 2X loading dye (without β-mercapto-ethanol and no heating of the sample) was loaded onto 10% polyacrylamide gels. Proteins were transferred onto nitrocellulose membranes using semi dry turbo transfer system. Then the membrane was incubated for overnight with primary antibodies diluted 1:1000 in 5 % BSA in TBST. Blots were washed with PBST (3X) before and after the secondary antibodies prior to imaging. Chemiluminescence images were obtained using ChemiDoc Imaging Systems (Bio-Rad) using west femto reagents.

### Cell viability studies

Reagents and protocol were purchased from CellTiter-Glo® Luminescent Cell Viability Assay (Promega). We conducted Cell-titre-glo (CTG) assays on the AML cell lines by adding 20µL of CTG reagent to 100µL of cells culture media before being placed on an orbital shaker for 2 minutes and subsequently incubated at room temperature for 10 minutes. Cell viability was measured using Hidex Sense microplate reader.

### Caspase3/7 glow

Cells were seeded in 384-well plates with 6,000 cells/well in 30uL of RPMI media and drug. After incubation, 30uL/well of Caspase-Glo 3/7^®^ reagent was added before being placed on an orbital shaker to equilibrate for 2 hours. Luminescence readings were measured using Hidex Sense microplate reader.

### LDH release assay

Reagents were purchased from CyQUANT™ LDH Cytotoxicity Assay (Thermo Fisher). AML cells (wild type, empty vector, and *DFNA5* OE) were seeded in triplicates using 96-well plates, at 8000 cells per well in 100μL OPTI-MEM media and drug. After incubation with the drug, 25ul was aliquoted into a new plate and added 25ul reaction mixture to each well and kept on shaker for 30min incubation. After 25ul of stop, solution was added to each well and kept in shaker for 5 min. Finally, we measured the OD values at 490nm and 680nm and LDH activity was measured by subtracting 680nm-490nm. The maximum LDH release was calculated with this formula.

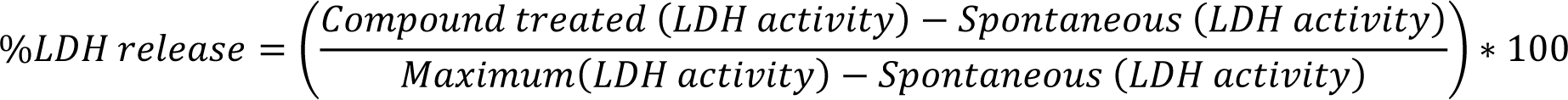

### Brightfield imaging

Cell was seeded in 5*10^5 cells/well in a 6 well plate in 2mL of RPMI media and drug. Then we treated the cells with azacytidine (1µM), venetoclax (1nM), azacytidine(1µM)+venetoclax(1nM) and etoposide (1uM). At the end of the incubation, the cells were transferred to a poly-D lysine coated 24 well plate and we stained with PI dye (1µM) into each well and imaged at 100X in brightfield imaging for ballooning morphology using Olympus FV1000 Confocal Microscope

### SYTOX green and Propidium Iodide (PI) uptake assay

Cells were seeded in 6-well plates with 500,000 cells/well in 2mL of RPMI media and drug. Subsequently, cells were stained with (i) 1µg/mL of PI dye for the PI uptake assay (ii) 1:3750 dilution of SYTOX green for the SYTOX green uptake assay. The number of PI positive or SYTOX green positive cells were imaged using Incucyte Live-Cell Analysis Systems (Sartorius) for every 30min to 4hrs.

### Coculture assay

EV cells and *DFNA5* KO (mCherry positive) cells were seeded in the ratio of 80:20 (total 10^5cells per well) respectively in a 96 well plate and treated with varying concentration of azacytidine, venetoclax and combination of venetoclax at 1nM and azacytidine at different doses. On day 3 and day 5 of the treatment FACS analysis was done to obtain the percentage of live cells present in the EV and *DFNA5* KO compartment after every alternate day of treatment with the drugs.

## Results

### Azacytidine upregulates *DFNA5* mRNA and protein levels via hypomethylation

Cell death caused by most chemotherapeutic drugs or targeted therapy drugs is mediated by activation of mitochondrial pathway of apoptosis [35]. Previous studies showed that pyroptosis, a non-apoptotic, gasdermin mediated form of programmed cell death, could also play a pivotal role in drug-induced cell death. However, switch in cell death mechanisms from apoptosis to pyroptosis in response to therapy depends on the expression level of *DFNA5*, pore forming protein [24]. We hypothesize that azacytidine as a hypomethylating agent can increase *DFNA5* expression in AML cells by demethylating *DFNA5* promoter. We performed methylation-specific PCR to measure cytosine methylation in *DFNA5* promoter region. To this end we treated MOLM-13 cells with 0.5µM azacytidine for up to 8 days and found significant reduction in the methylated promoter of *DFNA5* at day 6, 8 compared to day 0. **(Figure 1A)**.

**Figure 1.**
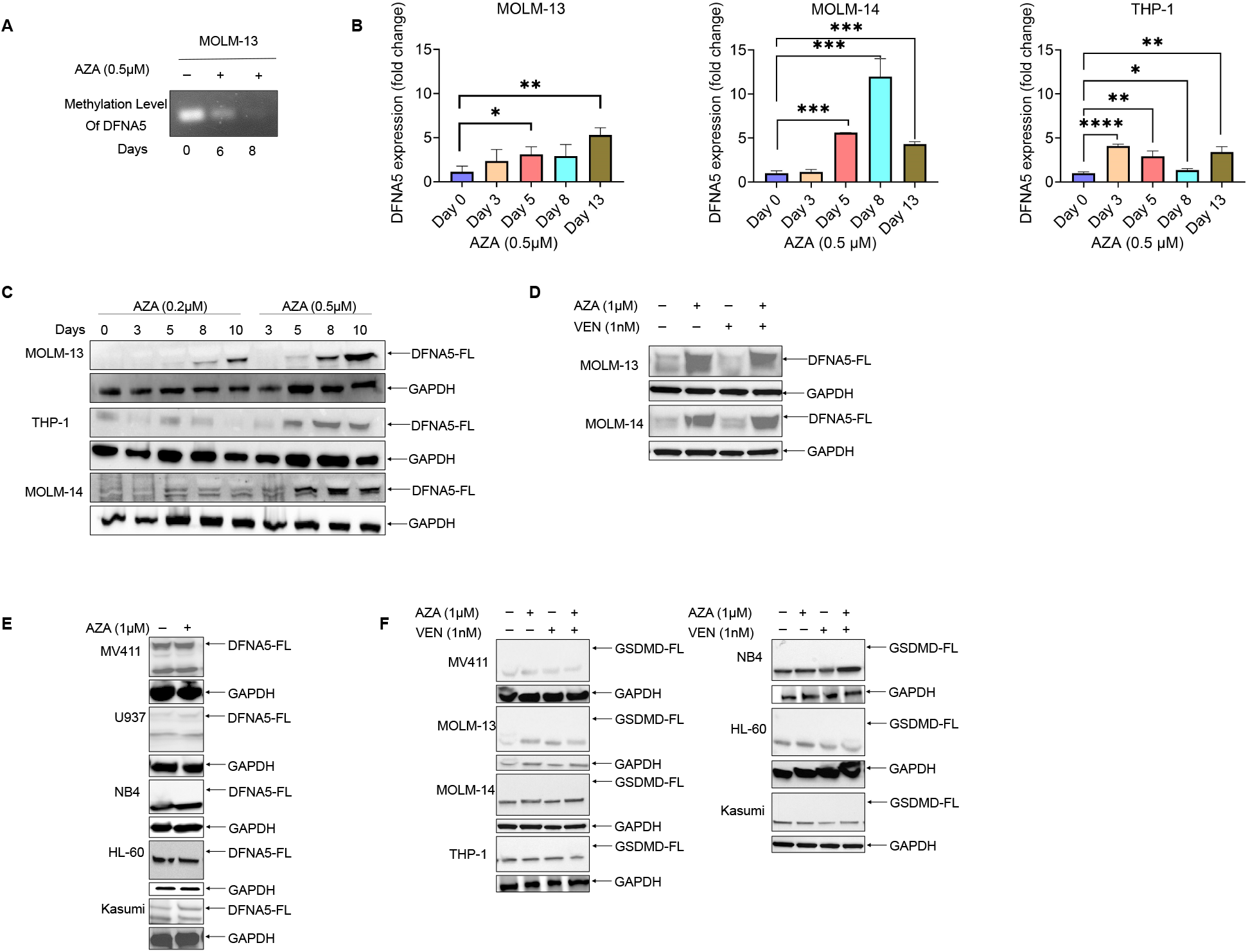
Azacytidine upregulates DFNA5 mRNA and protein levels via hypomethylation. **(A)** DNA methylation of the promoter region measured by methylation specific PCR (MSP) using methylated primers sets designed for promoter region of the DFNA5 gene in MOLM-13 cell lines treated with AZA at 0.5µM concentration. **(B)** Quantification of DFNA5 mRNA expression using real-time PCR following AZA treatment at 0.5µM for every alternate day up to 13 days in MOLM-13, MOLM-14 and THP-1 cell lines. Error bars indicate mean± SD, n=3; *P<0.05, **P<0.01, ***P<0.001, two-tailed Student’s t-test. **(C, D)** Immunoblotting of DFNA5 upon treatment with different AZA concentrations **(C)** (0.2µM and 0.5µM) for every alternate day up to 10 days in MOLM-13, THP-1 and MOLM-14 cell lines. **(D)** AZA (1µM), VEN (1nM) and their combination for up to 72 hrs post treatment. **(E)** Immunoblotting of DFNA5 upon treatment with AZA (1µM) concentration for up to 72 hrs post treatment in MV411, U937, NB4, HL-60, and Kasumi cell lines. **(F)** Immunoblotting of GSDMD upon treatment with AZA (1µM) VEN (1nM) and their combination for up to 72 hrs post treatment in MV411, MOLM-13, MOLM-14, THP-1, NB4, HL-60 and Kasumi cell lines.

We next asked whether the demethylation could result in subsequent transcriptional increase in *DFNA5* mRNA. Using *DFNA5*-specific qPCR primers, we confirmed significant upregulation of *DFNA5* at mRNA levels across 3 AML cell lines (MOLM-13, MOLM-14 and THP-1) with 0.5µM azacytidine (**Figure 1B**). Of note, the time of induction of *DFNA5* levels was variable across cell lines. We then assessed *DFNA5* protein expression using consecutive treatment with azacytidine for 10 days. We observed upregulation in *DFNA5* protein from as early as 3 days, though there was heterogeneity in response amongst different cell lines (**Figure 1C**). In THP-1 and Molm-14, *DFNA5* expression increased after 5 days post azacytidine, while MOLM-13 showed upregulation after 8 days of azacytidine treatment. As expected, higher azacytidine concentration showed *DFNA5* upregulation as early as 3 days in MOLM-13 and MOLM-14 (**Figure 1D**). Of note, temporal differences in *DFNA5* protein levels correlated with *DFNA5* transcript levels **(Figure 1B and 1C)**.

We failed to observe *DFNA5* protein upregulation in MV411, U937, NB4, HL-60 and Kasumi or T-ALL line (Jurkat) even after 1µM of azacytidine treatment because different cell lines have different levels of methylation[36] **(Figure 1E, Supplementary** Figure 1A). Besides *DFNA5*, other members of gasdermin family such as *GSDMD, GSDMA*, *GSDMB* and *GSDMC* also possess pore-forming gasdermin-N domains. However, except for *GSDMD*, none of the other Gasdermins are shown to be cleaved in response to physiological or pathological stimuli. Hence, we next checked *GSDMD* upregulation after azacytidine treatment. We failed to see upregulation of *GSDMD* in MOLM-13, MOLM-14, THP-1, NB4, HL-60 and Kasumi cell lines (**Figure 1F**). This suggests that *DFNA5* but not *GSDMD* is hypomethylated by azacytidine and may contribute towards pore formation in azacytidine treated AML cells.

### Azacytidine cleaves *DFNA5*-FL to *DFNA5*-NT and oligomerises to cause membrane rupture in AML cell lines

Unlike pore-forming toxins, *DFNA5* is cytoplasmic protein that must assemble oligomers to penetrate the plasma membrane from inside to induce pyroptosis[25]. We sought to evaluate the functional relevance of azacytidine-mediated *DFNA5* upregulation. Full-length *DFNA5* does not possess cytotoxic activity and hence mere upregulation of *DFNA5* expression alone is insufficient to initiate pyroptosis. Rather, the truncated form due to cleavage of *DFNA5*-FL to *DFNA5*-NT results in pyroptosis[25]. To investigate whether *DFNA5* upregulated by azacytidine is subsequently cleaved to form functional *DFNA5*-NT, we treated THP-1 cells with 0.5µM azacytidine for up to 10 days. Using pooled protein lysates from both pellet and supernatant for immunoblotting and found that azacytidine induced *DFNA5* cleavage to form *DFNA5*-NT as early as 3 days post-treatment (**Figure 2A**). To validate that the *DFNA5*-NT fragment can form stable oligomers, we performed in vitro oligomerization assay followed by non-reducing polyacrylamide gel electrophoresis to detect higher molecular weight oligomers as well as dimers (70kD) (**Figure 2B**). To further distinguish whether azacytidine-mediated *DFNA5* cleavage and oligomerization is due to epigenetic effects (hypomethylation) or non-epigenetic effects, we ectopically expressed *DFNA5* two AML cell lines **(Figure 2C, 2G, Supplementary** Figure 1B). Overexpression of *DFNA5* in THP-1 cells showed robust *DFNA5*-NT cleavage with in 20hrs post azacytidine treatment (**Figure 2D**) and enhanced cell death in both THP-1 and MOLM-14 DFNA5 OE cells compared to EV cells **(Supplementary** Figure 1C**, 1D)**. *DFNA5* OE THP-1 and AML-2 cells showed robust oligomerization indicated by smear formation compared to EV cells in response to azacytidine (**Figure 2F, 2G**). This suggests that AML cells with abundant expression of *DFNA5* can activate N-terminal cleavage independent of hypomethylation effects of azacytidine. We predicted that the azacytidine mediated *DFNA5* cleavage is due to activation of intrinsic apoptosis pathway that causes caspase activation downstream of mitochondrial outer membrane permeabilization. Hence, we next turned to venetoclax and found that at lower doses (1nM), venetoclax failed to cause *DFNA5* cleavage in THP-1 *DFNA5* OE cells. However, at venetoclax at higher concentration (1uM) caused *DFNA5* cleavage in THP-1 *DFNA5* OE cells (**Figure 2D, 2E**). We suspect that the combination of azacytidine+venetoclax showed similar level of *DFNA5*-NT and oligomers as azacytidine monotherapy due to lack of venetoclax-mediated *DFNA5*-NT cleavage at lower concentration (**Figure 2E**). Thus, this data collectively suggests that azacytidine plays dual role of epigenetically upregulating endogenous *DFNA5* via hypomethylation as well as inducing direct cleavage of *DFNA5*-FL into *DFNA5*-NT.

**Figure 2.**
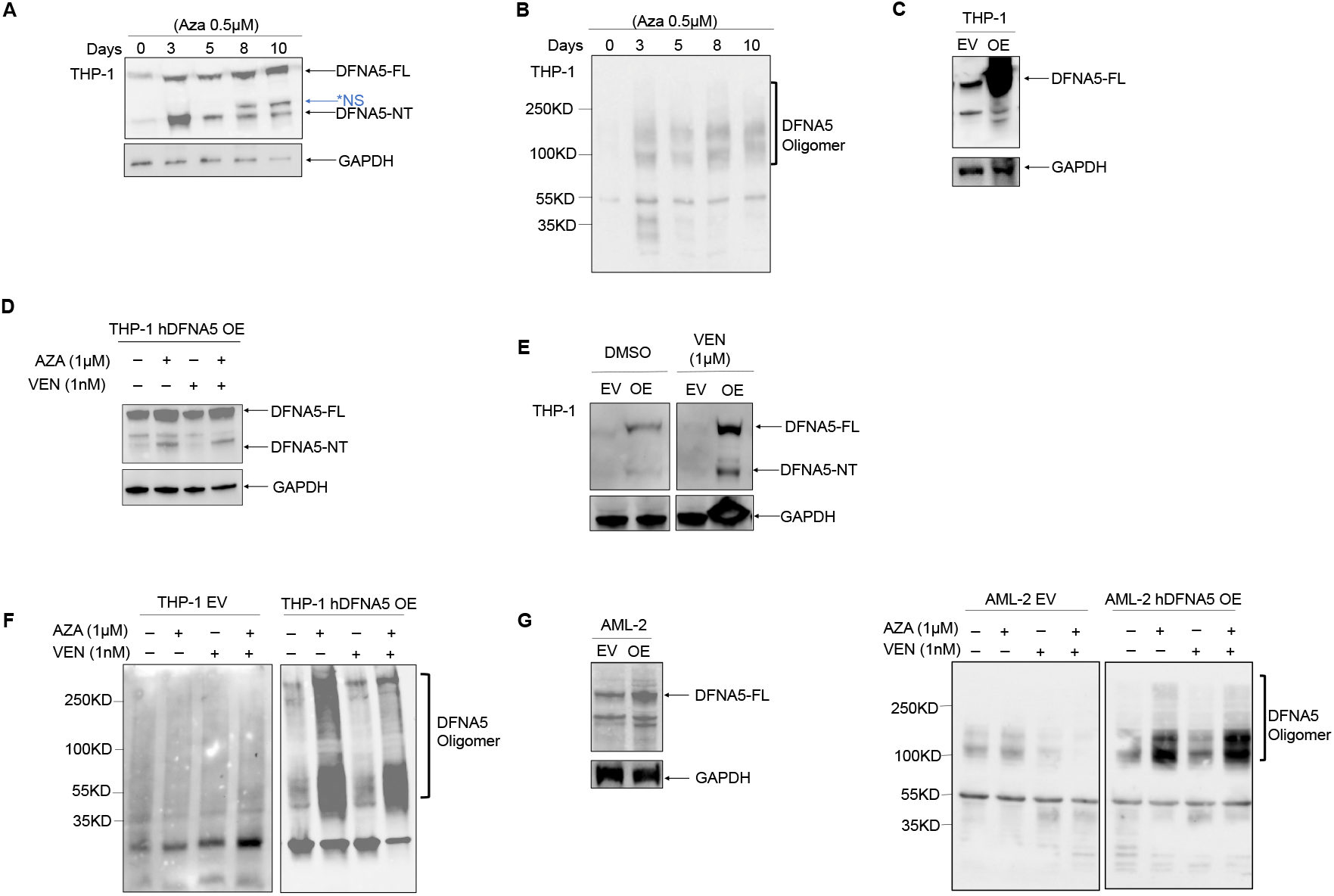
Azacytidine cleaves DFNA5 -FL to its functional DFNA5 –NT and oligomerises to cause membrane rupture in AML cell lines. **A)** Immunoblotting of DFNA5-NT protein upon long term treatment with AZA (0.5µM) in THP-1 cell lines. *NS=Non-Specific band **(B)** Non-reducing PAGE to measure DFNA5-NT oligomers upon long term treatment with AZA (0.5µM) in THP-1 cell lines. **(C)** Immunoblotting of hDFNA5 to confirm the over expression (OE) in THP-1 cell line post 1 week of puromycin selection. **(D, E)** Immunoblotting of DFNA5-NT in **(D)** THP-1 hDFNA5 OE cells upon treatment with AZA (1µM) and VEN (1nM) and its combination for 24hrs post treatment. **(E)** THP-1 EV and hDFNA5 OE cells upon treatment with high concentration of VEN (1µM) for 24hrs post treatment. **(F, G)** Non-reducing PAGE to measure DFNA5-NT forming oligomers upon treatment with AZA (1µM), VEN (1nM) and their combination for 24hrs in **(F)** THP-1 EV and hDFNA5 OE **(G)** AML-2 EV and hDFNA5 OE after confirmation of OE of hDFNA5 in AML-2 cell lines.

### Azacytidine induces membrane rupture and pyroptosis of AML cells

We so far showed that azacytidine increases *DFNA5* expression via hypomethylation and undergoes proteolytic cleavage to form *DFNA5*-NT and oligomerization. We next tested whether azacytidine-induced *DFNA5*-NT oligomers possess membrane-spanning pore forming action that results in osmotic swelling and plasma membrane rupture— characteristics of pyroptotic cell death. To test this, we measured lactate dehydrogenase (LDH) level, a marker for membrane rupture in 4 AML cell lines (THP-1, AML-2, AML-3 and MOLM-14) after treatment with venetoclax, azacytidine and azacytidine+venetoclax. We transduced 4 AML cell lines with *DFNA5* via CRISPR lentivector system to drive stable overexpression of *DFNA5*.

We measured LDH release in *DFNA5* OE cells instead of endogenous cells because azacytidine-mediated *DFNA5* upregulation requires long-term azacytidine stimulation in RPMI media (with serum), which leads to interferes with LDH signal due to colorimetric interactions. Azacytidine caused a statistically significant increase in LDH release in *DFNA5* OE THP-1 (P<0.01), AML-2 (P<0.01), AML-3 (P<0.01), Molm-14 (P<0.01) and Jurkat (P<0.001) cells compared to empty vector (EV) control (**Figure 3A-D**). Low dose venetoclax (1nM) alone or in combination with azacytidine caused minimal LDH release compared to high dose (1µM), as seen with oligomerization and DFNA-5 cleavage studies (**Figure 2E**).

**Figure 3.**
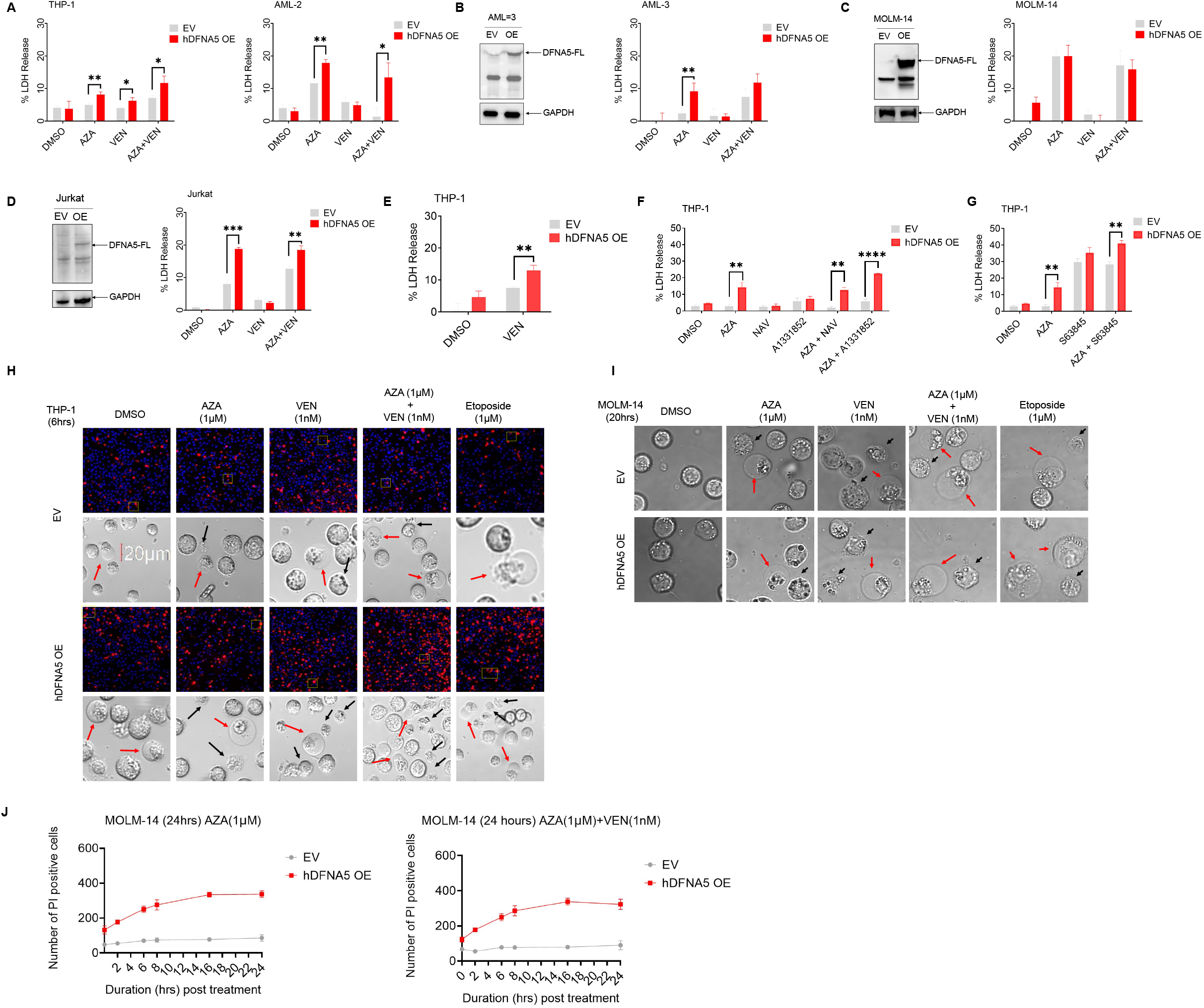
AML cells exhibit DFNA5 dependent cell cytotoxicity in response to venetoclax and azacytidine combination therapy. **(A-D)** LDH release measurement assay upon treatment with AZA (1µM), VEN (1nM) and their combination for 24hrs in (A) EV and hDFNA5 OE of THP-1 and AML-2 **(B)** AML-3 EV and hDFNA5 OE after confirmation of hDFNA5 OE by immunoblotting **(C)** MOLM-14 EV and hDFNA5 OE after confirmation of hDFNA5 OE by immunoblotting. **(D)** Jurkat EV and hDFNA5 OE after confirmation of hDFNA5 OE by immunoblotting **(E-F)** LDH release measurement assay upon treatment with **(E)** high concentration of VEN (1µM) for 24hrs. **(F)** BH3 mimetics Navitoclax (NAV)(1nM), A1331852 (100nM) and the combination with AZA (1µM) **(G)** S63845 (1µM) and combination with AZA (1µM) for 24hrs. Error bars indicate mean± SD, n=3 *P<0.05, **P<0.01, ***P<0.001, two-tailed Student’s t-test. **(H, I)** Brightfield images obtained during AZA (1µM), VEN (1nM) and its combination treatment **(H)** in THP-1 EV and hDFNA5 OE for 6hrs along with PI uptake **(I)** in MOLM-14 EV and hDFNA5 OE cells for 24hrs. **(J)** Uptake of PI quantified by live cell imaging in Incucyte microscope monitored for 24hrs post treatment with AZA (1µM), AZA(1µM)+VEN (1nM) and its combination in MOLM-14 EV and hDFNA5 OE. Etoposide (1µM) was used as positive control.

We next tested whether other BH3 mimetics drugs (besides venetoclax) that activate intrinsic apoptosis singling such as BCL-XL (A1331852), MCL-1 (S63845) and BCL-2/BCL-XL (navitoclax) inhibitors would also cause membrane rupture in combination with azacytidine. Except for navitoclax, both A1331852 (P<0.01) and S63845 (P<0.01) showed significant increase in LDH release in *DFNA5* OE cells compared to EV control when combined with azacytidine **(Figure 2F, G)**. Collectively, this data shows that short-term treatment with azacytidine can induce *DFNA5*-dependent membrane rupture indicated by LDH release. This explains that if *DFNA5* is expressed then azacytidine can cause direct cytotoxic effects via non-apoptotic forms of cell death. Of note, this effect is independent of hypomethylating effects of azacytidine as it is caused after short-term treatment.

### AML cells exhibit *DFNA5-*dependent pyroptosis in response to azacytidine and venetoclax combination therapy

After showing evidence for azacytidine-mediated membrane rupture, we next sought to determine whether *DFNA5*-dependent membrane rupture results in pyroptosis. Since pyroptotic cells become leaky and permeable after losing the membrane integrity, we measured propidium iodide (PI) uptake as a marker of pyroptosis. We measured PI uptake after treatment with azacytidine, venetoclax and the combination (azacytidine+venetoclax). *DFNA5* OE caused remarkable increase in PI uptake in response to azacytidine+venetoclax after 6 hours compared to DMSO or single agents (venetoclax or azacytidine) in THP-1 cells (**Figure 3H**). In addition, longer treatment exposure (24h) also caused increased PI uptake in THP-1 *DFNA5* OE cells upon treatment with azacytidine and azacytidine+venetoclax (**Supplementary** Figure 2C). Overall, these data establish that AML cells with higher *DFNA5* levels can cause increased PI uptake via azacytidine and venetoclax combination. Of note, apoptosis would also result in PI uptake and hence venetoclax treatment resulted in higher PI uptake compared to DMSO which was further increased in *DNFA5* OE cells (**Figure 3H** **and Supplemental Figure 2**. We further quantified PI uptake by measuring PI^+^ cells using Incucyte. Compared to EV, *DFNA5* OE cells showed increased PI uptake in azacytidine, and combination treated MOLM-14 cells (**Figure 3J**).

Since cells undergoing other forms of necrosis, apart from pyroptosis, can also exhibit membrane rupture (indicative via LDH release and PI uptake), we next gathered morphological evidence to confirm pyroptosis induced by azacytidine or azacytidine+venetoclax. The characteristic feature of pyroptotic cells is “ballooning-like” morphology —cell swelling with characteristically large bubbles from cell membrane. We utilized brightfield imaging to visualize cell morphology of *DFNA5* OE THP-1 and MOLM-14 cells after treatment with azacytidine, venetoclax, azacytidine+venetoclax. In comparison to EV cells, *DFNA5* OE cells displayed cellular shrinkage and classical membrane ballooning morphology (fried egg structures) at 24-hour post treatment with azacytidine, venetoclax and azacytidine+venetoclax combination **(Figure 3H, 3I and Supplemental Figure 2A-C)**. As previously shown *DFNA5* OE resulted in pyroptosis of etoposide treated cells used as positive control here.[24]

Next, we asked about the onset of pyroptosis in response to treatment, given that pyroptosis is a rapid phenomenon. We performed time course imaging at 2, 4 and 6 hours post treatment in THP-1 cells with *DFNA5* OE or EV. We observed a time dependent increase in the number of pyroptotic cells from 2 – 6 hours with azacytidine+venetoclax combination **(Supplement Figure 2A-D)**. In summary, we found that ectopic expression of *DFNA5* sensitizes AML cells towards azacytidine+venetoclax induced pyroptosis.

### *DFNA5* deletion protects AML cells from azacytidine+venetoclax mediated pyroptosis

While OE of *DFNA5* resulted in increased pyroptosis, we also observed evidence of apoptotic cells. Hence to determine direct contribution of *DFNA5* in mediating total cell death, we knocked out (KO) *DFNA5* using CRISPR-Cas-9 system. To validate the KO, we exposed MOLM-14 KO cells with azacytidine (0.5µM) for 8-days. We observed upregulation of *DFNA5* expression in EV cell but not in KO cell, confirming the KO of *DFNA5* gene was successful (**Figure 4A**).

**Figure 4.**
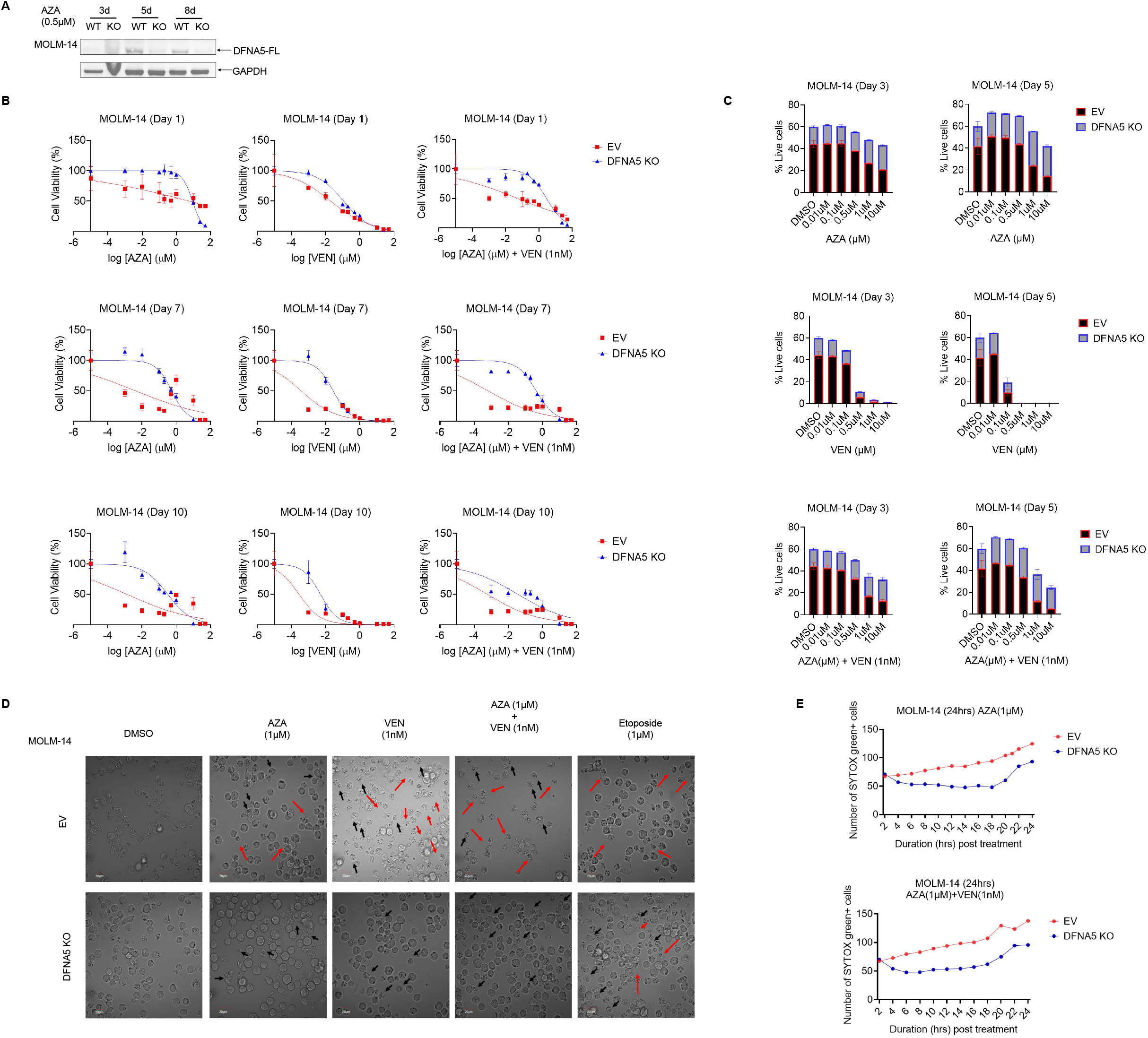
AML cells undergo DFNA5 dependent cell death on azacytidine, venetoclax and combination treatment. **(A)** Immunoblotting of DFNA5 to confirm the KO in MOLM-14 cells treated with AZA (0.5uM) for alternate days up to 8 days. **(B)** Cell viability was assessed at day 1, 7 and 10 for MOLM-14 EV and DFNA5 KO cell lines upon treatment with varying concentration of AZA, VEN and combination of VEN at 1nM and varying concentrations of AZA using cell titre glow (CTG) assay. Error bars: Mean±SEM, n=3 replicates **(C)** Similar concentrations of VEN, AZA and combination were used to perform co-culture assay (with 20% KO cells and 80% EV cells) and the % live cell readings were obtained at day 3 and 5 in FACS Error bars: Mean±SEM, n=3 replicates. **(D)** Brightfield images obtained during AZA (1µM), VEN (1nM) and its combination treatment in MOLM-14 EV and DFNA5 KO cells for 24hrs. **(E)** Uptake of SYTOX green quantified by live cell imaging in Incucyte microscope monitored for 24hrs post treatment with AZA (1µM), VEN (1nM) and its combination in MOLM-14 EV and DFNA5 KO cells Error bars indicate mean± SD, n=6.

Having seen that *DFNA5* OE resulted in pyroptosis and apoptosis after treatment with azacytidine, we first quantified the total cell viability in response to azacytidine, venetoclax and azacytidine+venetoclax using cell titre glow (CTG) reagent and coculture assay. We treated MOLM-14 EV and *DNFA5* KO cells with increasing concentrations of azacytidine, venetoclax and azacytidine+venetoclax combination at every alternate day for over 10 days. *DFNA5* KO cells showed increased viability compared to EV MOLM-14 after treatment with azacytidine, venetoclax and azacytidine+venetoclax combination **(Figure 4B and 4C)**. At day 7, we found that *DFNA5* KO increased IC50 of azacytidine (IC50 DNFAKO-0.5478 µM vs EV IC50-0.004192 µM), venetoclax (IC50 DNFAKO-28.01 nM vs EV IC50-0.3693 nM) and the combination (IC50 DNFAKO-434 nM vs EV IC50-0.6514 nM). Compared to venetoclax, azacytidine mediated cytotoxicity is more dependent on *DFNA5*. To further confirm that DNFA5 is required for cytotoxicity induced by azacytidine and venetoclax, we next measured total cell count by flowcytometry after co-culture of EV (80%) and *DFNA5* KO (20%) MOLM-14 cells in the presence of azacytidine, venetoclax and the combination. However, at day 3 of co-culture, the total number of live cells is 60% as we eliminated the debris, dead cells, and doublet cell and gated on live cell population based on forward scatter X side scatter. Interestingly, we observed that MOLM-14 EV cells showed dose-dependent reduction after days 3 and 5 post treatment, while *DFNA5* KO cells remained viable at day 3 even at 10uM azacytidine and started to proliferate by day 5. This further confirms that DNFA5 is required for azacytidine-mediated cell killing in MOLM-14. While *DFNA5* KO protected MOLM-14 cells from venetoclax-mediated killing at 100nM, higher doses of venetoclax were able to eradicate MOLM-14 cells irrespective of DFNA-5 KO. Similar to azacytidine alone, combination of azacytidine+venetoclax showed selective sensitivity against EV cells as opposed to *DFNA5* KO cells. This suggests that *DFNA5* is required for azacytidine+venetoclax mediated cell killing. Through the brightfield imaging, we show that *DFNA5* KO cells underwent more apoptosis instead of pyroptosis compared to the EV, that showed less number of ballooning cells during the treatment (**Figure 4D**). To further confirm that *DFNA5* KO prevents azacytidine or azacytidine+venetoclax mediated pyroptosis, we carried out SYTOX green staining, marker for membrane rupture. Of note, we could not use PI uptake since KO cells were selected using m-Cherry fluorescence. We observed significant reduction in the SYTOX green positive cells after *DFNA5* KO cell compared to the EV cells over a 24h measurement induced by azacytidine (p<0.0001) and azacytidine+venetoclax (p<0.0001) (**Figure 4E**). Collectively, we found that azacytidine-mediated pyroptosis is *DFNA5* dependent and contributes to total cytotoxicity induced by azacytidine+venetoclax.

### Azacytidine-mediated *DFNA5* cleavage is caspase-3 dependent

Thus far, we found that azacytidine enhances *DFNA5* expression and cleaves *DFNA5*-FL to form *DFNA5*-NT oligomers which causes cell membrane rupture and subsequent pyroptosis. Wang et al. reported that the knockdown of caspase 3 prevented *DFNA5*-FL cleavage, which provided us insight that the *DFNA5* acts downstream of caspase 3 [26]. We next asked whether azacytidine-mediated *DFNA5* cleavage was direct or mediated via caspase-3 (**Figure 6A**) [16]. To test the direct role of azacytidine in mediating DFNA-5 cleavage, we performed qPCR and WB to check if the hypomethylating agent azacytidine upregulates caspase 3 expression in MOLM-13, MOLM-14 and THP-1 cell lines. We observed that the azacytidine did not affect the expression of caspase 3 at both mRNA and protein levels after long term treatment with azacytidine (0.5µM) **(Figure 5A, 5B)**. However, we observed increased levels of cleaved caspase 3 during the long-term treatment with azacytidine (0.5µM) at as early as day 3 in all the three AML cell lines (**Figure 5B**). This data suggests that azacytidine does not upregulate the caspase 3 levels but instead activates the caspase 3 cleavage upon treatment (**Figure 5C**). Next, we performed caspase 3/7 glow assay to quantify the cleaved caspase 3 levels after azacytidine treatment. We observed that MOLM-13, MOLM-14 and THP-1 cell lines showed significantly increased active caspase 3 after treatment with azacytidine, venetoclax and the combination compared to DMSO (**Figure 5D**).

**Figure 5.**
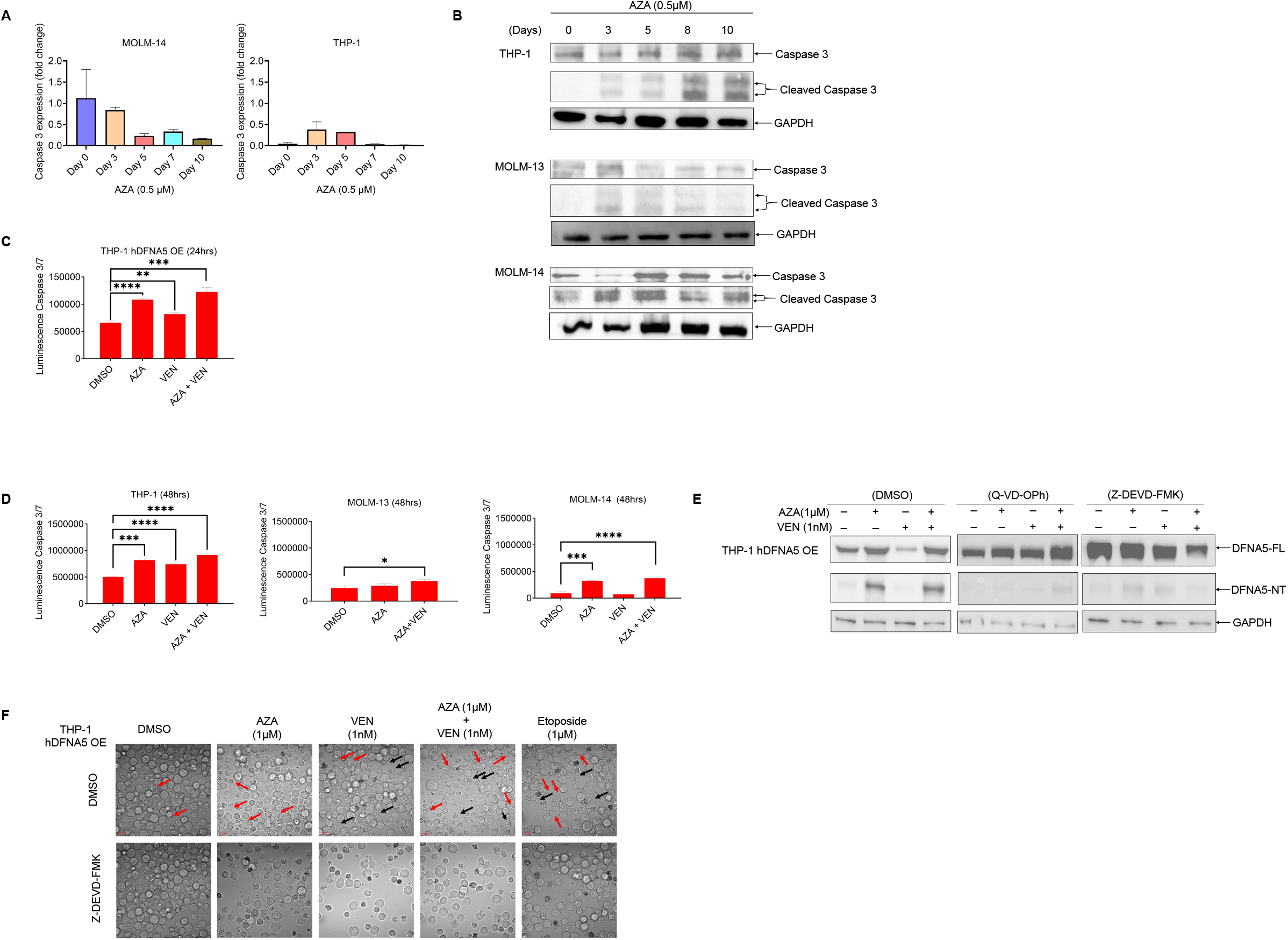
Azacytidine-mediated DFNA5 cleavage is caspase-3 dependent. **(A)** Quantification of mRNA expression of caspase-3 following AZA treatment at 0.5µM for every alternate day up to 10 days in MOLM-14 and THP-1 cell lines Error bars indicate mean± SD, n=3: two-tailed Student’s t-test. **(B)** Immunoblotting of caspase-3 and cleaved caspase-3 in ThP-1, MOLM-13, MOLM-14 after 0.5µM AZA treatment for every alternate day up to 10 days. **(C, D)** Quantifying caspase 3/7 activation by measuring the luminescence with Caspase3/7 Glo assay in **(C)** THP-1 hDFNA5 OE cells treated with AZA (1uM), VEN (1nM) and the combination for 24 hrs. **(D)** MOLM-13, MOLM-14 and THP-1 treated with AZA (1µM), VEN (1nM) and the combination for 48 hrs. Error bars indicate mean± SD, n=3 *P<0.05, **P<0.01, ***P<0.001, two-tailed Student’s t-test. **(E)** Immunoblotting of DFNA5-NT in THP-1 hDFNA5 OE cells after caspase inhibition by Q-VD-OPH and caspase-3 inhibition by Z-DEVD-FMK followed by 24hrs treatment with AZA (1µM), VEN (1nM) and its combination. **(F)** Brightfield images obtained after caspase-3 inhibition with Z-DEVD-FMK and upon treatment with AZA (1µM), VEN (1nM) and its combination in THP-1 hDFNA5 OE cells for 24hrs. Etoposide (1µM) was used as positive control.

**Figure 6.**
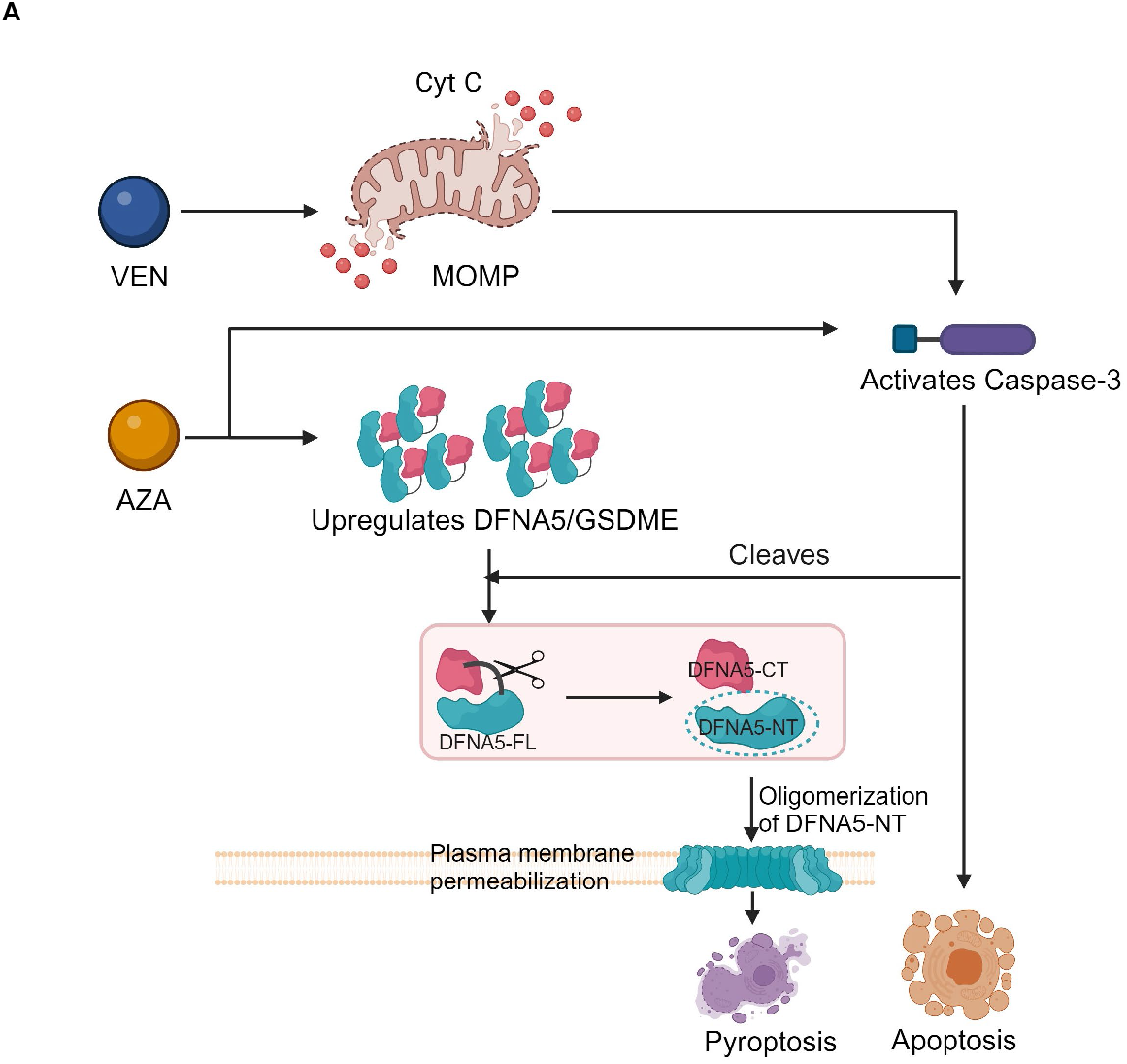
**(A)** Schematic diagram showing the proposed role of AZA in both upregulating DFNA5 and mediating DFNA5 cleavage by activating Caspase 3 to trigger pyroptosis by itself and in combination with VEN at higher concentration. This represents the role of upregulated DFNA5 in AML cell and causing enhanced cell death during the combination therapy.

Next, we sought to verify the direct contribution of caspase 3 in cleaving *DFNA5*. To test this, we first inhibited THP-1 *DFNA5* OE cells with the pan-caspase inhibitor (Q-VD-OPh) and caspase-3 specific inhibitor (Z-DEVD-FMK) for 4 hours before subjecting these cells to azacytidine, venetoclax and azacytidine+venetoclax for 24 hours. Immunoblotting of DFNA-5 showed that pretreatment with pan-caspase inhibitor as well as caspase-3 inhibitor prevented *DFNA5* cleavage (**Figure 5E**). Finally, we performed brightfield imaging and confirmed that the blockade of caspse-3 using Z-DEVD-FMK prevented azacytidine, venetoclax or the combination induced pyroptosis and apoptosis (**Figure 5F**). Collectively, these results suggest that the azacytidine activates the caspase 3 and is an upstream modulator in *DFNA5* mediated pyroptosis in AML cells.

## Discussion

Despite advancements in AML therapy, high relapse rates among patients call for improvements in current treatments. With increasing literature on pyroptosis as an alternative cell death mechanism, we sought to understand whether azacytidine+venetoclax therapies exerted cytotoxic effects via non-apoptotic pathways.

Our study explored the interactions between azacytidine and *DFNA5* as well as its consequent activation of different cell death pathways in AML cell lines MOLM-13, MOLM-14, THP-1 and AML-2. We used various techniques (CTG, Co-culture, PI Uptake assay and Brightfield Imaging) to analyse the cell death pathways induced by azacytidine, venetoclax and azacytidine+venetoclax.

We initially hypothesised that venetoclax synergises with azacytidine by activating caspase-3, which then cleaves *DFNA5* to form *DFNA5*-NT, a common pyroptosis mediator. However, our results suggest that azacytidine on its own can induce pyroptosis of AML cells via epigenetic mechanisms after long-term treatment. We also provide an explanation for why AML cells do not die via pyroptosis in response to treatment with chemotherapy or targeted therapy. This is due to hypermethylation of tumor suppressive genes like *DFNA5*. Thus, methylation of the *DFNA5* promoter region silences *DFNA5* expression and hence, prevents pyroptosis. Our study for the first time showed the mechanism by which clinically relevant doses of azacytidine upregulates endogenous *DFNA5* expression levels in AML cell lines: THP-1, MOLM-13 and MOLM-14.

Additionally, it appears that some AML cell lines are much more sensitised towards pyroptosis by azacytidine (MOLM-13, MOLM-14, THP-1, AML-2, and AML-3) compared to other cell lines, like Kasumi, NB4, HL-60 and MV411. Simultaneously, our data suggest azacytidine itself can cleave caspase-3 to mediate pyroptosis. We found that azacytidine plays 2 significant roles in mediating pyroptosis: 1) it hypomethylates human *DFNA5* to upregulate *DFNA5* mRNA and protein levels and 2) it can cleave and activates caspase 3, which in turn causes *DFNA5* cleavage and oligomerization to induce pyroptosis of AML cell lines.

While we primarily explored azacytidine’s mechanism in driving pyroptosis, we also found that higher concentration of venetoclax treatment can trigger pyroptosis and with evidence suggesting that venetoclax can induce pyroptosis through BCL-2/*DFNA5* pathway in myeloid cells [37], it was worthy to explore pyroptosis induced during the synergistic killing via azacytidine and venetoclax.

With the cause of AML relapse being tumour regrowth stemming from chemotherapy resistant, apoptosis-evading cells, we see a dire need to explore other cell death pathways to circumvent relapses. Our results provide evidence that pyroptosis is one of such pathways which can be tapped on and needs to be further validated clinically.

Through our research, we have gained a much better understanding of the role that azacytidine plays in upregulating *DFNA5* expression, as well as inducing pyroptosis in AML cell lines. We remain optimistic that our findings would build on existing research to refine the understanding of the role that non-apoptotic mechanisms play in AML treatment. This could eventually be applied to augment the efficacy of current treatment methods and significantly improve the quality of care we provide our patients with.

