## Supplementary Figures for "*DFNA5*-mediated pyroptosis is a driver for venetoclax and azacytidine synergy in myeloid leukemia"

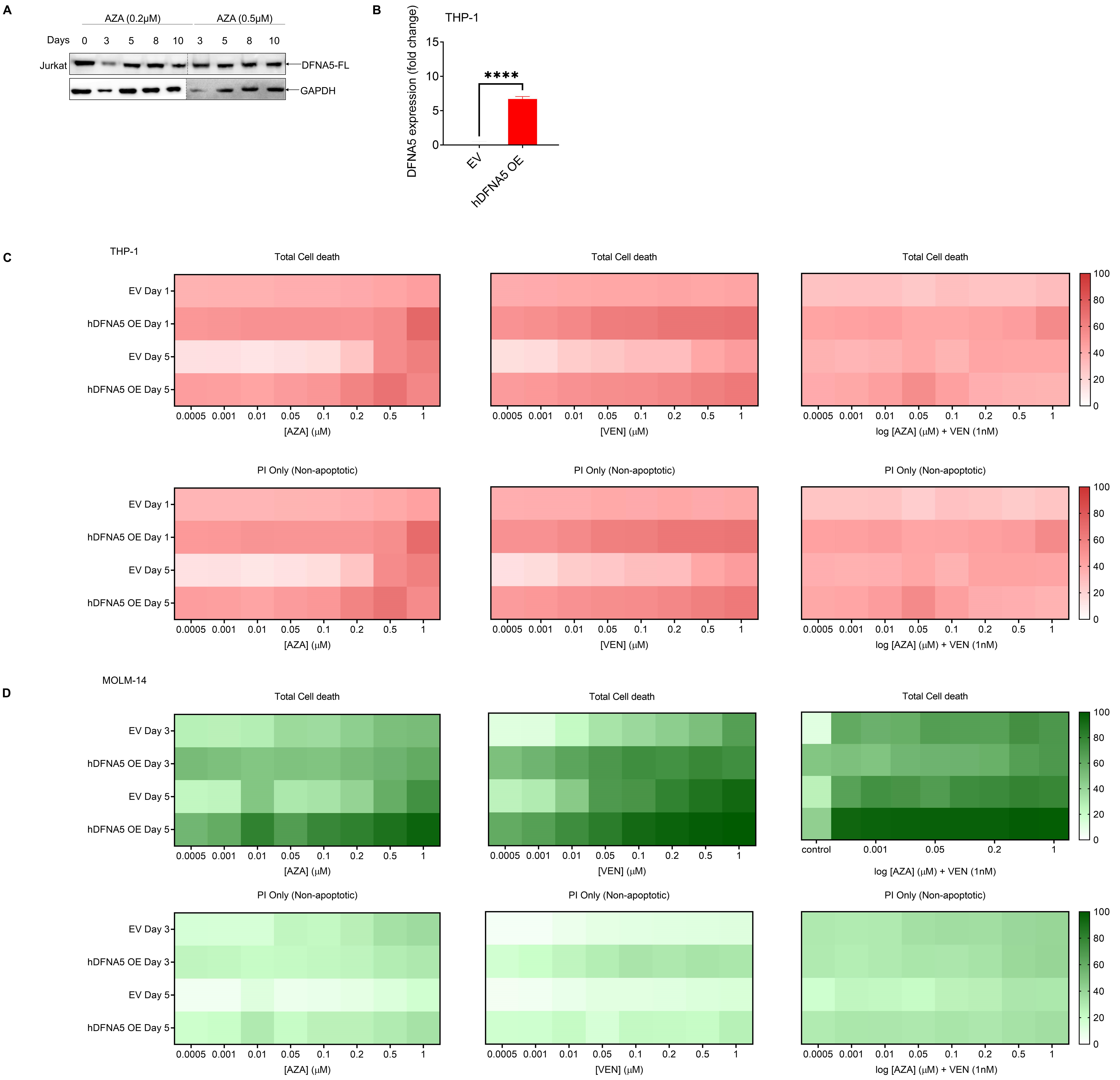


**Supplementary Figure 1. (A)** Immunoblotting of DFNA5 upon treatment with AZA at 0.2µM and 0.5µM concentration in Jurkat cell line **(B)** Quantification of mRNA expression of DFNA5 to confirm the OE in THP-1 cell line Error bars indicate mean± SD, n=3; two-tailed Student's t-test. **(C, D)** Cell death type was analyzed using Annexin V-PI assay performed by FACS to analyze the total cell death and PI only positive graph for **(C)** THP-1 EV and hDFNA5 OE cells. **(D)** MOLM-14 EV and hDFNA5 OE cells. The experiment was performed in three biological replicates.

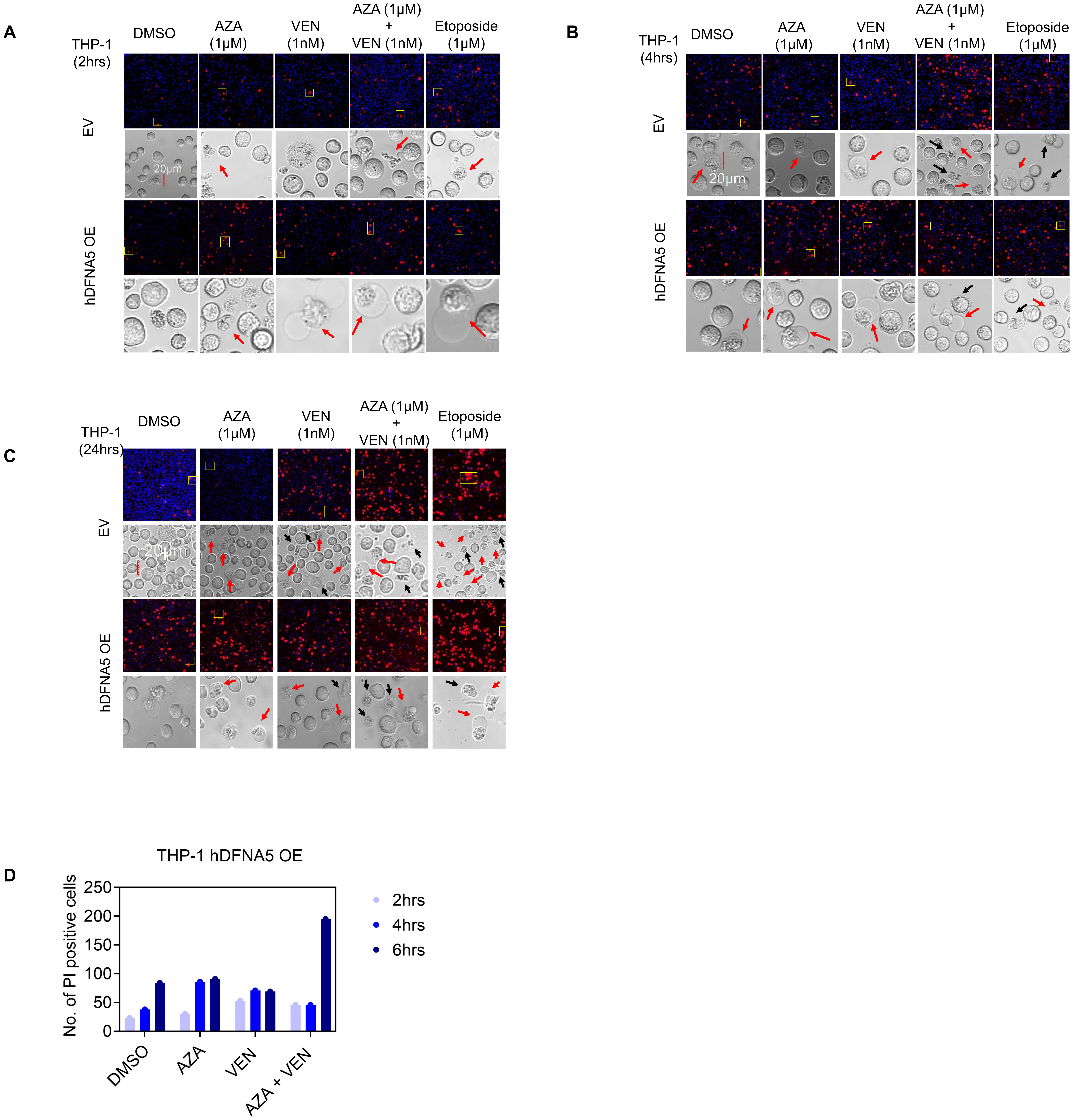


**Supplementary Figure 2. (A, B, C)** Quantification of uptake of PI and brightfield images of THP-1 EV and hDFNA5 OE cells at **(A)** 2 hrs **(B)** 4hrs and **(C)** 24hrs **(D)** Quantification of PI uptake of the cells during the treatment of AZA (1μM), VEN (1nM) and its combination.
